# Essential role of hsa-miR-203a-3p in type I Interferons immune homeostasis during Influenza and NDV infection

**DOI:** 10.1101/2025.04.01.646562

**Authors:** Pramod kumar, Ashish Kumar, Akhilesh Kumar, Himanshu Kumar

## Abstract

MicroRNAs (miRNAs) are small, non-coding molecules that act as essential post-transcriptional regulators in various biological processes. Many studies suggest that miRNAs may modulate the host’s immune response during virus infections. We analyzed publicly available transcriptomics data involving infection with different RNA viruses and identified the most prominent candidate, miR-203a-3p. This miRNA is upregulated during H7N9, HCV, and DENV2 infections. Interestingly, pathway analysis of microRNA-targeted genes shows that miR-203a-3p targets multiple genes of the type-I interferon and JAK-STAT pathway. In this study, we reported the novel role of miR-203a-3p as it is elevated in response to polyinosinic-polycytidylic acid [poly(I:C)] transfection and infection with various RNA viruses such as Newcastle Disease Virus (NDV) and A/PR8/H1N1 influenza virus. We found that miR-203a-3p promotes the A/PR8/H1N1 virus replication by suppressing the host’s type-I interferons and interferon-stimulated genes. We demonstrated that the miR-203a-3p overexpression reduced the expression of ISGs and is attributed to the direct binding of miR-203a-3p to 3’ UTRs of Janus-activated kinase 1 (JAK1), STAT1, SOCS3, and multiple IFNA transcripts. Altogether, these findings strongly suggest that miR-203a-3p plays a pivotal role in immune homeostasis by regulation of type I IFN and supports A/PR8/H1N1 and NDV infection by targeting multiple genes of the host’s type I IFNs and JAK-STAT signaling pathways.

## Introduction

RNA viruses like influenza, MERS, SARS-CoV, and SARS-CoV-2 have been the cause of several epidemics and pandemics in the recent past and, therefore, pose a significant threat to global human health. These viruses are first sensed by the host’s innate immune system, which consists of a family of extracellular and intracellular sensors (1, 2). The innate immune sensors consist of various families of pattern recognition receptors (PRRs) that sense the conserved molecular signatures associated with viruses, such as viral protein, RNA with 5′-triphosphate ends, hypo- or unmethylated viral DNA, single-stranded RNA (ssRNA), and double-stranded RNA (dsRNA) (3, 4) . Sensing of PAMPs by PRRs activates a plethora of downstream signaling pathways, culminating in the activation of transcription factor interferon (IFN) regulatory factor (IRFs) and nuclear factor-kB (NF-kB) for the induction of Type-I interferons and pro-inflammatory cytokines(5, 6) . All these factors collectively develop an antiviral state. Type-I interferons further establish antiviral effects through the induction of an array of genes called interferon-stimulated genes (ISGs) (7). This complex process of antiviral signaling is regulated at the post-transcriptional and post-translational levels. Post-translational regulation primarily occurs through the ubiquitination or phosphorylation of transcription factors such as IRFs, JAKs, and STATs(8–10) . In contrast, post-transcriptional regulation is largely mediated by various non-coding RNAs, including long non-coding RNAs (lncRNAs), circular RNAs (circRNAs), and microRNAs (miRNAs) (11–13). The miRNAs, a class of single-stranded non-coding RNAs (ncRNAs) ranging from 18 to 21 nucleotides in length, play a pivotal role in fine-tuning antiviral immune responses using a multiprotein RNA-induced silencing complex(14).

The miRNAs regulate gene expression by binding to the 3’UTR (3’ untranslated region) region of target transcripts and repressing their translation. Several transcriptomic studies have shown that the virus infection induces the expression of many miRNAs, targets innate immune signaling pathways, and modulates the antiviral response (15, 16). Notably, we or other groups demonstrated that the miRNAs can also directly bind to the genomic RNA of RNA viruses and affect viral replication (17–21). Upon infection, the genetic material of RNA viruses is sensed by Toll-like receptor 3 (TLR3), TLR7, and TLR8 in the endosomal compartment; it is also sensed by the cytosolic sensors such as RIG-I and MDA5 in the cytosol-inducing type-I interferons necessary to mount an effective antiviral immune response(22–24).

The type-I interferons consist of several members of IFNα and a single IFN-β and act in a paracrine and autocrine manner on various immune and non-immune cells through interferon receptors consisting of IFNAR1 and IFNAR2. Upon binding of interferon to the receptor, the cascade of downstream signaling is activated by JAK1 and tyrosine kinase 2 (TYK2), which in turn activate STAT1 and STAT2 and form an IFN-stimulated gene factor 3 (ISGF3). Not only does STAT1/2 form a heterodimer, but STAT1/STAT1 also forms a homodimer, also known as IFNγ activating factor (GAF), where ISGF3 and GAF bind to the ISREs and IFNγ activation sites leading to transcription of numerous ISGs, whose integrated action results in an antiviral state(25–28).

This study identified that miR-203a-3p is induced upon RNA virus infection and poly(I:C) transfection. The miR-203a-3p targets the 3’UTR of the IFNA and IFNA signaling pathway gene members (IFNAR1, IFNA2, IFNA4, IFNA7, IFNA10, IFNA14, IFNA16, IFNA17, IFNA21) through RISC. Ectopic expression of miR-203a-3p in HEK-293T, A549, and HeLa cells diminishes the type-I interferon-mediated antiviral response, thereby increasing the viral load.

## Material and Methods

### miRNA Identification and pathway analysis

NCBI-GEO database was queried for miRNA profiling experiments and databases relevant to H7N9, HCV, and DENV2 infections. GSE150782, GSE40744, and GSE135311 were selected based on experimental design and relevance to the study. Differentially expressed miRNAs were identified in control and infected samples of the given datasets using the GEO2R online tool (29, 30). TargetScan 7.1 (31, 32) and miRabel (33) were used as default parameters to predict the target genes of selected miRNAs.

### Cells, transfection, plasmids, mimics, and inhibitors

A549 human adenocarcinoma alveolar basal epithelial cells (Cell Repository, NCCS, India), HeLa cervical cancer cells (Cell Repository, NCCS, India), and HEK293T human embryonic kidney cells (ATCC, CRL-3216) were cultured in Dulbecco’s modified Eagle’s medium (DMEM) supplemented with 10% fetal bovine serum (FBS) and 1% antibiotic-antimycotic solution (Gibco). Cells were maintained at 37°C in a humidified incubator with 5% CO_2_.

For transfection experiments, cells were seeded in 6, 12, or 24-well cell culture plates depending on experiment requirements. Transfections were carried out when cells reached 70- 90% confluency. Plasmid, miRNA mimics (Invitrogen), inhibitors, and negative control or poly I:C were transfected according to the manufacturer’s protocol in HEK293T and HeLa cells using Lipofectamine 2000 (Invitrogen), and Lipofectamine 3000 (Invitrogen) was used for A549 cells, and respective samples were collected at desired time points.

### Viruses and infection

Influenza virus (strain PR8, A/PR8/H1N1) and GFP-tagged NDV (strain LaSota) viruses were used for infection experiments. Viral stocks were prepared in pathogen-free embryonated chicken eggs and stored at -80, The multiplicity of infection (MOI) was calculated based on viral titer and cell density(34, 35).

Cells were seeded in 6 or 12 well tissue culture plates at densities of 0.3 x 10^6^ or 0.1 x 10^6^ cells/well. 24 hours post-transfection cells were washed with SF-DMEM or 1x Phosphate-buffered saline (PBS) and infected with NDV-GFP at 0.5 MOI and A/PR8/H1N1 at 0.5, 1, and 2 MOI. Virus suspensions in SF-DMEM were added to the cells and incubated for 1 hour at 37. Cells were then washed with SF-DMEM or 1x Phosphate-buffered saline (PBS) incubated in complete DMEM and supplemented with 1% FBS. Replacement media for A/PR8/H1N1 infection was supplemented with L-1-Tosylamide-2-phenylethyl chloromethyl ketone (TPCK)-treated trypsin (1 μg/ml). GFP-tagged NDV was kindly provided by Dr. Peter Palese, Icahn School of Medicine at Mount Sinai, New York, USA.

### Luciferase reporter assay

Cells were seeded in 24-well plates and, after 24 hours, transfected with 150 ng/well UTR-luc- construct and 320 ng/well of EV/p203 plasmid. PRLTK was used as a control at a 20 ng/well concentration. After 24 hours, cells were lysed using a passive lysis buffer provided with the Dual-Glo Luciferase reagent assay kit.

Firefly luciferase luminescence was measured by mixing 50 µl of Dual-Glo Luciferase Reagent with 50 µl of cell lysate in an opaque 96-well plate. To measure the Renilla luciferase activity, an equal volume of Stop & Glow reagent was added per well. After 10 minutes of incubation, the luminescence of Renilla luciferase was measured. The luminescence intensity of renilla luciferase was used to normalize the firefly luciferase luminescence; Luminescence signals were quantified using Promega GloMax multi-microplate reader

### Quantitative real-time reverse transcription-PCR

Total RNA was extracted using TRIzol reagent (Invitrogen) and converted into cDNA using the iScript cDNA synthesis kit (Bio-Rad) following the manufacturer’s protocol. quantitative reverse transcription polymerase chain reaction (qRT-PCR) was performed using gene-specific primers (sequences provided in supplementary table) using CYBR green chemistry (Bio-Rad) on the QuantStudio 3 Real-Time PCR system; 18S rRNA was used as an internal control for normalization. TaqMan-based qRT-PCR was conducted for miRNA expression analysis using a universal PCR master mix (Applied Biosystems) and specific TaqMan probes for U6 and miR-203a-3p. U6 served as the internal control for normalization.

### Fluorescence-activated cell sorting (FACS) analysis

Cells were seeded in six-well plates and, at 60–70% confluency, transfected with pMIR-203a-3p or pMIR-empty vector as control. After 24 hours, cells were washed with phosphate-buffered saline (PBS) and infected with NDV-GFP at 1.5 MOI for 1 hour. Infected cells were washed with PBS, replenished with 1% FBS containing DMEM, and incubated overnight. Cells were detached using a 0.25% trypsin-EDTA solution, washed with PBS, and fixed with 4% paraformaldehyde for 5 minutes. Fixed cells were washed twice with PBS resuspended in PBS and analyzed using a BD FACSAria II flow cytometer. FlowJo software (BD Biosciences) was used to analyze the data, and the fluorescence intensity was measured to quantify the levels of GFP expression.

### RNA immunoprecipitation

RNA immunoprecipitation was performed as described in the previous articles (36, 37). HEK 293T cells were lysed in lysis buffer (0.5% NP-40, 150 mM KCl, 25 mM tris-glycine (pH 7.5) and incubated with M2 Flag affinity beads overnight at 4°C. Beads were washed 5 times with washing buffer (300 mM NaCl, 50 mM Tris-glycine [pH 7.5], 5 mM MgCl, 0.05% NP-40). Following the manufacturer’s protocol, Trizol reagent was used to get RNA from immunoprecipitated ribonucleoproteins (RNPs). The quality and quantity of extracted RNA were assessed using a NanoDrop spectrophotometer.

### Measurement of viral titer

Madin-Darbin Canine Kidney (MDCK) cells were used for viral titer assays. Cells were infected with A/PR8/H1N1 or NDV in the serum-free MEM medium with viral supernatant. After 1 hour of virus infection, cells were washed with 1X PBS and were overlaid with 1.2% Avicel (Sigma-Aldrich; equivalent to Avicel RC-581 from FMC Corp.) in 2 ml maintenance medium and incubated at 37°C with 5% CO for 96 hours. Avicel overlays were removed, and the cells were fixed with 4% paraformaldehyde for 20 min. Fixed cells were stained with a 0.5% crystal violet solution, and plaques were counted under the microscope using transmitted light microscopy. For the 50% tissue culture infective dose (TCID_50_) assay, infected monolayers were fixed and stained with 1% crystal violet after 96 h of infection, and the TCID_50_ was calculated based on the Reed and Muench method as described previously (38, 39).

## Results

### miR-203a-3p is induced during RNA virus infection

The miRNAs are key post-transcriptional regulators for gene expression, including genes involved in innate immunity. To find novel miRNA regulating antiviral innate immunity, we compared publicly available microarray data sets (GSE150782, GSE40744), and NGS (GSE135311) where different cells were infected with H7N9, HCV, and DENV2. Dysregulated miRNAs were identified based on a p-value threshold of <0.05 and a log2 fold change greater than 0.5 or less than -0.5. Among them, miR-203a-3p, miR-378c, and miR-199a-5p were the only miRNAs significantly upregulated across all three datasets (Fig. 1A). These three miRNAs were then analyzed for their target genes using TargetScan 7.2 and miRabel. The predicted targets were further examined through BioPlanet, KEGG pathway, and Reactome analyses to understand the affected pathways (40–43). Our findings indicate that miR-203a-3p plays a key role in regulating the Interferon alpha (IFNA) signaling pathway, TRAF6-mediated IRF7 activation, and the RIG-I-like receptor signaling pathway (Fig. 1B-C), suggesting this miRNA is crucial during RNA virus infections. These pathways were selected based on top-ranked p-values and odds ratios (Fig. S1A-B), highlighting their strong association with the identified gene set. As miR-203a-3p targets antiviral pathways and is upregulated during RNA virus infection (Fig. 1D-F), we decided to further study its role during RNA virus infection. To this end, first, we analyzed the expression of miRNA-203a-3p in PBMC, A549, HEK293 and HeLa cells upon A/PR8/H1N1 or NDV infection and found that these viruses induced the expression of miR-203a-3p multifold (Fig. 1G-K). Similarly, we observed the induction of miR-203a-3p upon transfection of A549 cells with poly(I:C), which activates RLR pathways (Fig. S1C). In contrast to miR-203a-3p, miRNA-378c and miR-199a-5p did not appear to target innate immune pathways (Fig. S2A–E). Collectively, these results suggest that RNA virus infection and mimicking RNA virus infection induces miR-203a-3p.

**Figure 1:**
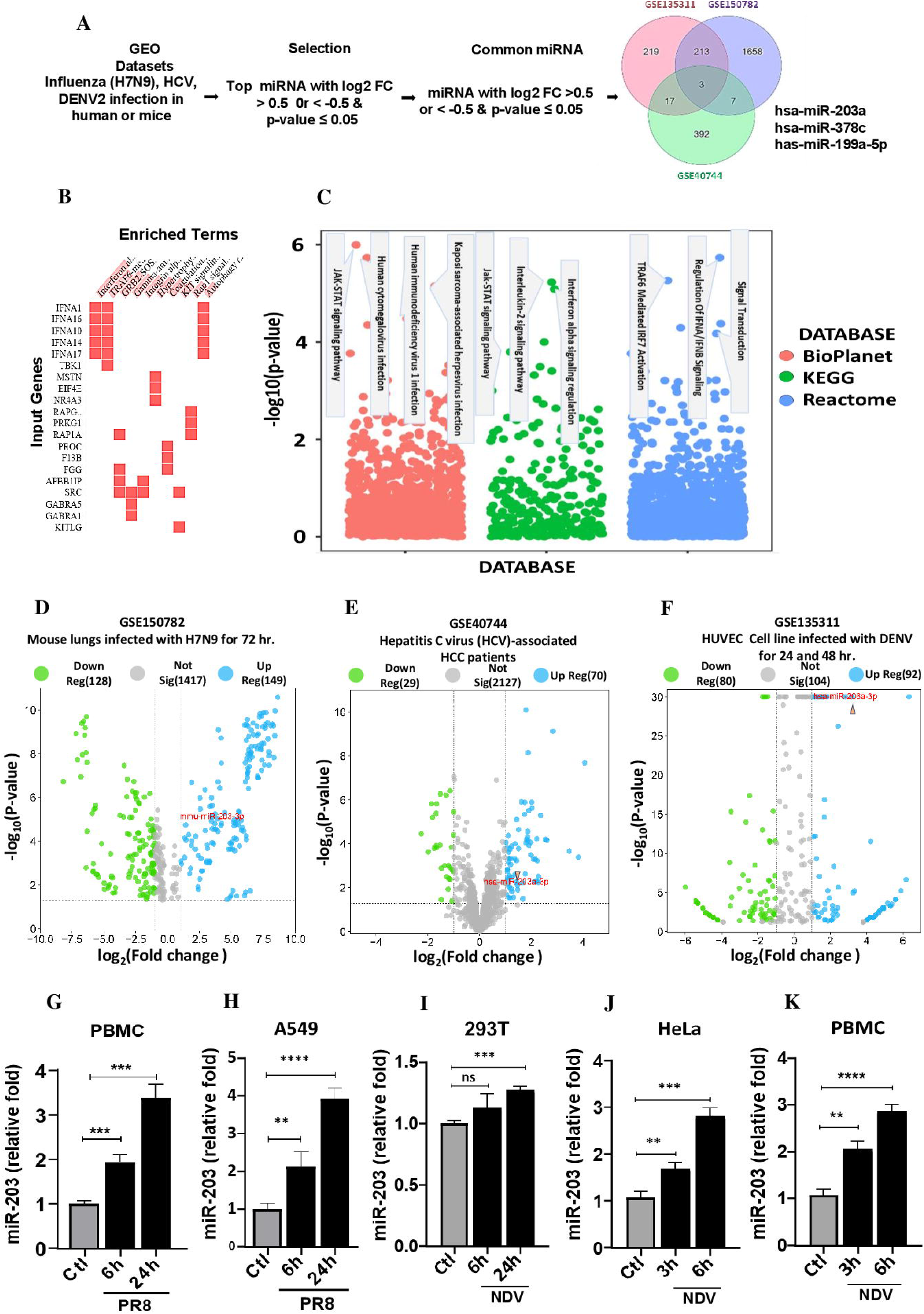
Identification and pathway analysis of miR-203-3p: (A) Microarray datasets for RNA virus infections, including H7N9, HCV and DENV2 were obtained from the Gene Expression Omnibus (GEO) database and analyzed using the GEO2R tool to identify differentially expressed miRNAs. Dysregulated miRNAs were selected based on a p-value < 0.05 and a log2 fold change (FC) threshold of >0.5 or < -0.5. The analysis identified miR-miR-203a-3p, miR-378c and miR-199a-5p were significantly upregulated across all three datasets. (B) Predicted target genes were analyzed using Enrichr to identify significantly enriched pathways. A clustergram visualizes genes involved in the top-enriched pathways based on Reactome pathway analysis. (C) Pathway enrichment analysis highlights the most significant pathways associated with the miRNA target genes across multiple databases, including BioPlanet, KEGG, and Reactome. (D-F) Volcano plots depict the differential expression of miR-203a-3p in the analyzed datasets. (G-K) Induction of miR-203a-3p in PBMCs, and HEK293, HeLa, and A549 cell lines at indicated time by qRT-PCR after infection with A/PR8/H1N1 and NDV. Statistical significance is indicated as follows: * (p < 0.05), ** (p < 0.01), *** (p < 0.001), and **** (p < 0.0001).

### miR-203a-3p directly targets the 3’UTR of IFNA members and the IFNA signaling pathway

In silico analysis indicates that miR-203a-3p binds to the 3’UTR of the interferon signaling pathway genes such as IFNAR1, JAK1, and STAT1, including various sub-members of interferon alpha (IFNA1, IFNA2, IFNA4, IFNA7, IFNA10, IFNA14, IFNA16, IFNA17) (Fig. 2A). IFNA is a predominant member of type-I interferon and a key contributor to the antiviral response. To investigate the IFNA members targeting by miR-203a-3p, we cloned full-length UTRs of these genes downstream to the luciferase reporter gene in the pMIR-REPORT vector. HEK293T cells were co-transfected with UTR-cloned luciferase reporter plasmids along with pmir-203a-3p (p203a) or an empty vector. Cells were lysed and subjected to luciferase assay after 48 hours of transfection. As expected, overexpression of miR-203a-3p significantly reduced the luciferase activity of IFNAR1, IFNA1, 4, 7, and 16, JAK1, STAT1, and SOCS3 (Fig. 2B-I), indicating that miR-203a-3p binds to 3’UTR of these genes, resulting in the downregulation of luciferase activity. However, luciferase activity did not change when IRF7 UTR lacking miR-203a-3p target sequence was co-transfected with miR203-3p (Fig. S3A).

**Figure 2:**
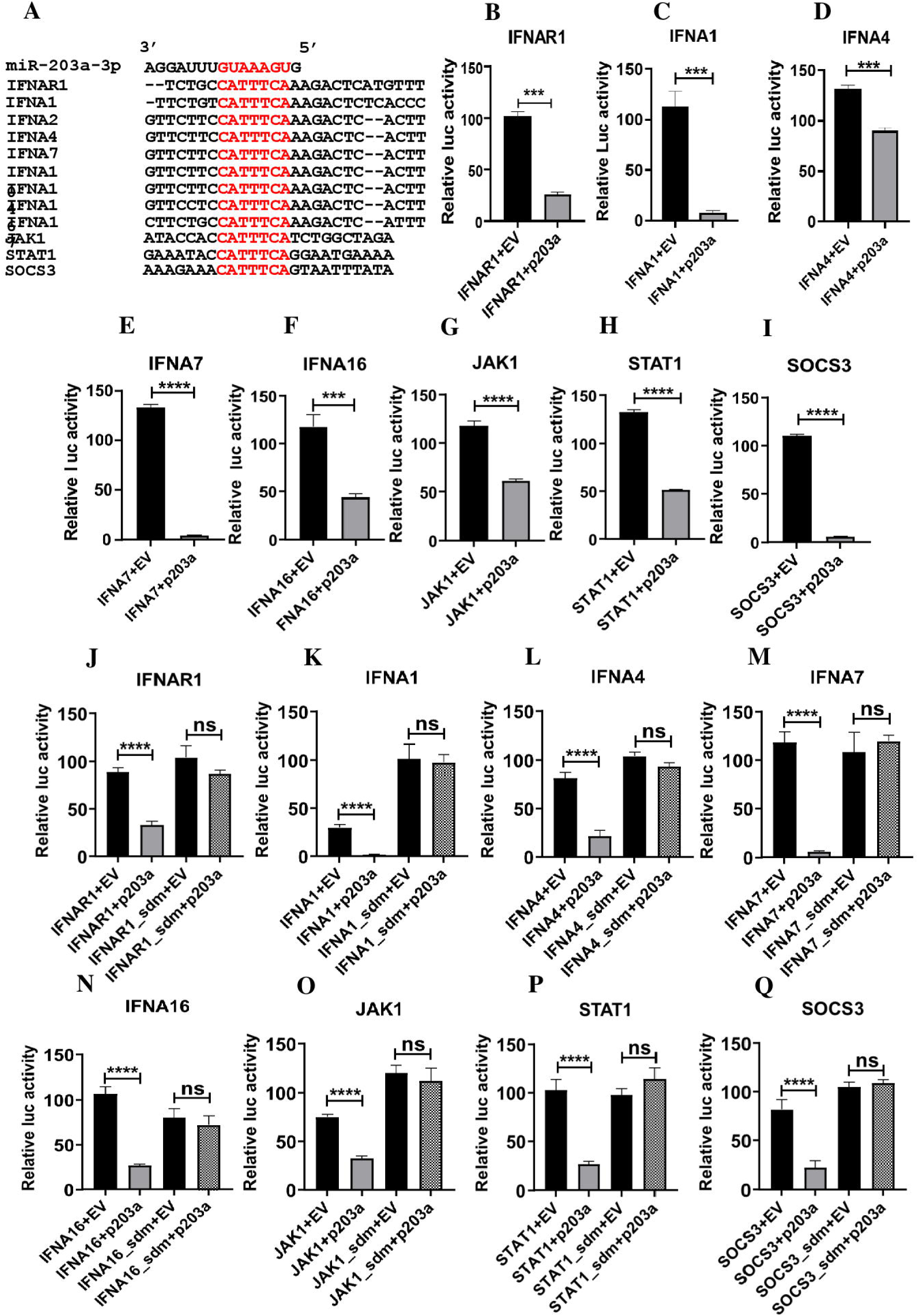
miR-203a-3p directly targets the 3’ UTR of IFNA members and it’s signaling pathway genes: (A) Schematic representation of conserved region in genes that are predicted to be targeted by miR-203a-3p, in the various members of type I interferon (IFN) and type I IFN signaling pathway. (B-I) The 3′ UTRs of IFNAR1, IFNA1, IFNA4, IFNA7, IFNA16, JAK1, STAT1, and SOCS3, were cloned into the pMIR-Report luciferase vector and co-transfected with either pMIR-203a-3p (p203) or an empty control vector. HEK293T cells were transfected with plasmids containing IFNAR1-Luc, IFNA1-Luc, IFNA4-Luc, IFNA7-Luc, IFNA16-Luc, JAK1-Luc, STAT1-Luc, or SOCS3-Luc, along with 50 ng of pRL-TK (Renilla luciferase for normalization), 100 ng of full-length wild-type (WT) 3′ UTR, and either 350 ng of the miR-203a-3p expression plasmid or 20 nM of a miR-203a-3p mimic. After 48 hours, cells were lysed, and luciferase activity was measured. (J-Q) The miR-203a-3p binding sites in the IFNAR1, IFNA1, IFNA4, IFNA7, IFNA16, JAK1, STAT1, and SOCS3 3′ UTRs were mutated using site-directed mutagenesis (SDM). The miRNA binding site sequence CATTTCA was mutated to ACTTGAC, and luciferase assay was performed in similar method as above as indicated constructs. The level of significance is indicated as follows: * (p < 0.01), ** (p < 0.001), *** (p < 0.0001), and **** (p < 0.00001). Assay. Statistical

To further validate the miR-203a-3p targeting sites in the 3’UTRs of target genes, the binding site were mutated from CATTTCA to ACTTGAC in the 3’ UTRs of the IFNAR1, IFNA1, IFNA4, IFNA7, IFNA16, JAK1, STAT1, and SOCS3 genes using Site-Directed Mutagenesis (SDM). After altering the miRNA binding site, the luciferase assay was performed as described previously. Luciferase assays with mutated constructs showed comparable luciferase activity with empty vector upon miR-203a-3p mimic transfection, confirming the specificity of miR-203a-3p’s interaction with the 3’UTR of target genes (Fig. 2J-Q).

Together, these results suggest that miR-203a-3p directly binds to the 3’UTRs of IFNA subtypes, IFNAR1, JAK1, STAT1, and SOCS3, leading to suppression of target gene transcript and may result to the downregulation type I IFN and type I IFN signaling pathway.

### miR-203a-3p target genes through RISC

MicroRNA target the transcripts through multiprotein complex known as RNA-induced Silencing Complex (RISC) and Ago2 is a one of essential component of RISC. To determine how miR-203a-3p regulates targeted transcripts. RNA immunoprecipitation (RIP) assay was performed along with Ago2. HEK293T cells were transfected with FLAG-tagged Ago2 and miR-203a-3p mimic (m203) or negative control (NC), followed NDV or A/PR8/H1N1 infection (Fig. 3A). Twenty-four hours post-infection, RNA associated with Ago2 was immunoprecipitated using FLAG antibody. Subsequently, the levels of Pan-α, IFNAR1, IFNA1, JAK1, and STATI transcripts were measured. The expression of these transcripts was significantly enriched in samples where miR-203a-3p were introduced compared to the negative control (NC). The overexpression of miR-203a-3p in cell was confirmed as shown in Fig. 3B. These results indicate the direct interactions between miR-203a-3p and Pan-α, IFNAR1, IFNA1, JAK1, and STATI mRNA transcripts. (Fig. 3C-G). Here, IRF7 was used as a non-target control (Fig. S3B).

**Figure 3:**
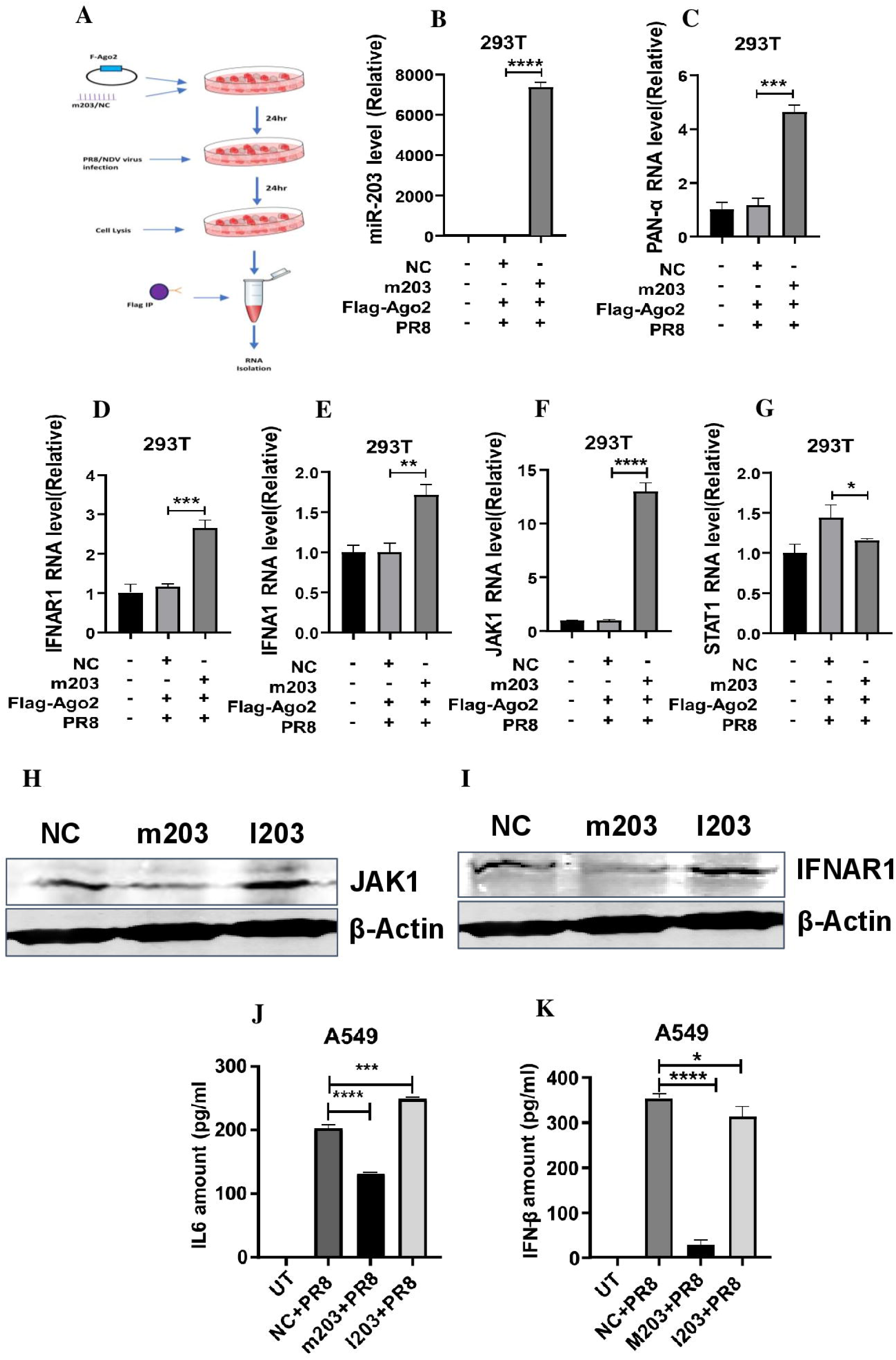
miR-203a-3p binds with Type I IFN pathway genes through Ago2-mediated RISC: (A) Schematic representation of the RNA immunoprecipitation (RIP) assay. HEK293T cells were either left untransfected or transfected with a *Flag-Ago2* plasmid along with either a negative control (NC) or *miR-203a-3p* (m203) mimic. Following transfection, cells were infected with A/PR8/H1N1 or NDV at an MOI of 1.5 for 24 hours. Cell lysates were subjected to RIP using an anti-Flag antibody. (B) The level of *miR-203a-3p* associated with the RNA-induced silencing complex (RISC) was quantified. (C-G) The mRNA levels of *pan-IFNA, IFNAR1, IFNA1, JAK1*, and *STAT1* in immunoprecipitated samples were measured using qRT-PCR. (H-I) A549 cells were transfected with NC, m203, or an inhibitor of miR-203a-3p (I203), followed by A/PR8/H1N1 infection. After 24 hours, cells were trypsinized, and protein lysates were collected for Western blot analysis. The protein levels of JAK1 and IFNAR1 were assessed using anti-JAK1 and anti-IFNAR1 antibodies. (J-K) A549 cells were transfected with NC, m203, or I203. After 24 hours, cells were infected with A/PR8/H1N1, and supernatants were collected 24 hours post-infection. and measured the IL-6 and IFN-β levels. by ELISA. Statistical significance is indicated as follows: * (p < 0.05), ** (p < 0.01), *** (p < 0.001), and **** (p < 0.0001).

Furthermore, these observations were confirmed at protein levels by western blot analysis using JAK1 or IFNAR1-specific antibodies. As both JAK1 or IFNAR1, play pivotal role in enhancement of type I IFN production in positive feedback loop fashion. As demonstrated above in RIP assay, the relative enrichment of these transcript for these genes were high. Therefore, we analyzed the protein of JAK1 and IFNAR1. As shown in the Fig, 3H and I, miR-203a-3p mimic (m203) significantly reduced the protein of IFNAR1 and JAK1 compared to the NC-transfected cells, whereas treatment with the miR-203a-3p inhibitor which sequester endogenous miR-203a-3p, modestly enhanced the expression of both IFNAR1 and JAK1 (Fig. 3H-I). Additionally, the effect of miR-203a-3p on IL-6 and IFN-β production in the culture supernatant was measured using ELISA. Notably, IL-6 production is driven by NF-κB, which is influenced by type I interferons. Similarly, IFN-β production is regulated through a positive feedback loop involving both IFN-α and IFN-β. To this end, A549 cells were transfected with NC, miR-203a-3p mimic (m203), or miR-203a-3p inhibitor (I203), followed by infection with A/PR8/H1N1 after 24 hours; the supernatant was collected 24 hours post-infection, and IL-6 protein levels were quantified using ELISA. As shown in Fig. 3J, the production of IL-6 was significantly reduced compared to the NC, whereas I203 transfection modestly enhanced the production of IL-6. Next, IFN-β production was also measured by ELISA. Cells transfected with miR-203a-3p showed marked reduction of IFN-β compared to the NC-transfected cells whereas introduction of I203 showed comparable levels of IFN-β with NC-transfected cells. (Fig. 3K), suggesting a broader regulatory role of miR-203a-3p in cytokine responses. Altogether, these results indicate that miR-203a-3p directly interacts with IFNA family genes and IFNAR1-JAK1-STAT1 signaling axis and suppresses at both the RNA and protein levels, thereby modulating type I IFN signaling.

To further validate the functional relevance of IFNAR1-JAK1-STAT1 signaling axis, a type-I IFNs non-responsive cells, Vero-E6 were infected with A/PR8/H1N1 or NDV at a multiplicity of infection (MOI) of 1.5. Viral load was assessed using qRT-PCR at 12 and 24 hours post-infection, showing an increase in viral load over time (Fig. S4 A-B). These results indicate that miR-203a-3p directly interacts with the transcripts of type I IFN members and type I IFN signaling molecules, including JAK1 and IFNA subtype genes, through RISC and reduce the production of type I IFN.

### miR-203a-3p suppresses antiviral responses

Viral infections are known to trigger the upregulation of innate immune cytokines, particularly, type-I interferons such as IFN-α and IFN-β, which are essential for activating interferon-stimulating genes (ISGs) that collectively inhibit viral replication and enhance immune responses. To assess and validate the impact of miR-203a-3p on type-I interferon (IFN) expression, A549 cells were transfected with negative control (NC), miR-203a-3p mimic (m203), or miR-203a-3p inhibitor (I203), followed by infection with A/PR8/H1N1 or NDV. We observed that miR-203a-3p mimic significantly reduced IFN-α and IFN-β transcript at various time points post-infection with both A/PR8/H1N1 and NDV in A549 cells and hPBMCs (Fig. 4A-D). Conversely, treatment with the miR-203a-3p inhibitor (I203) restored IFN-α and IFN-β expression (Fig. 4A–C). To investigate the downstream effect of reduced expression of Type I interferons (IFN-α and IFN-β), we analyzed the activation of the Interferon Stimulated Response Element (ISRE), a promoter element critical for ISG activation. Using an ISRE luciferase reporter (pISRE-luc) plasmid containing ISRE promoter present in the ISGs. The ISRE sequence cloned upstream of a luciferase gene, HEK293T cells were transfected with either a negative control (NC) or m203a-3p, along with pISRE-Luc. After 24 hours, the cells were either left untreated or treated with IFN-β for 24 hours. The results revealed a significant reduction in luciferase activity in cells transfected with m203a-3p whereas, luciferase activity was comparable in cell transfected with ISRE and ISRE along with NC (Fig. 4E). These findings suggest that miR-203a-3p suppresses Type I IFN signaling, resulting in reduced expression of IFN-α and IFN-β and subsequently ISG.

**Figure 4:**
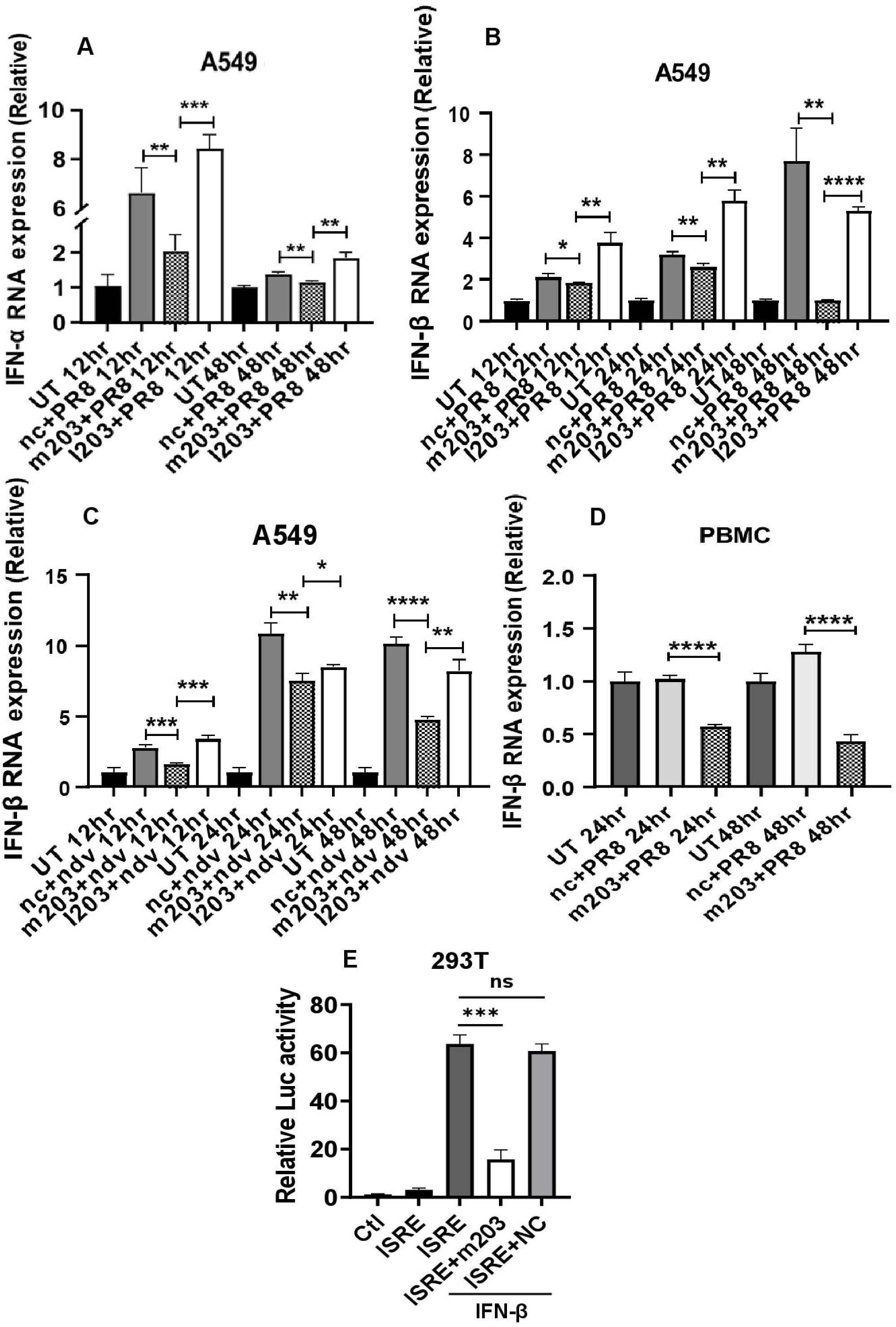
miR-203a-3p Overexpression suppresses type-I IFN and type-I IFN-inducible responses: (A) A549 cells were transfected with a miR-203a-3p mimic (m203a), a negative control (NC), or a miR-203a-3p inhibitor (I203) at a concentration of 50 nM. After 24 hours, cells were infected with A/PR8/H1N1 (MOI = 1.5). Samples were collected at 12 and 48 hours post-infection to measure IFNα expression by qRT-PCR. (B–C) Similarly, IFN-β expression was measured at 12, 24, and 48 hours following A/PR8/H1N1 or NDV infection. (D) The level of IFN-β was also measured in peripheral blood mononuclear cells (PBMCs),which were transfected with m203a or NC (50 nM), followed by A/PR8/H1N1 infection. The relative expression of IFN-β was measured by qRT-PCR at 24, and 48 hours post-infection. (E) HEK293T cells were transfected with either pISRE alone, pISRE + m20a, or pISRE + negative control (NC). After 24 hours, the cells were stimulated for 24 hours with IFN-β, and luciferase activity was measured. Statistical significance is indicated as follows: * (p < 0.05), ** (p < 0.01), *** (p < 0.001), and **** (p < 0.0001).

### miR-203a-3p promotes the RNA virus replication

To determine the effect of miR-203a-3p on viral load, A549 cells were transfected with negative control (NC), miR-203a-3p mimic (m203), or m203 inhibitor (I203). Subsequently, the transfected cells were infected with A/PR8/H1N1 or Newcastle Disease Virus (NDV) at a multiplicity of infection (MOI) of 1.5. The viral load was then quantified at 12, 24, and 48hours post-infection by quantitative PCR, expression of the A/PR8/H1N1 NP gene (for influenza), and the NDV L-pol gene, which encodes the large polymerase of NDV, was measured. A significant increase in viral load was observed in cells transfected with the miR-203a-3p mimic (m203) compared to the negative control, while the viral load was reduced in cells treated with the miR-203a-3p inhibitor (I203) (Fig. 5A-B). To further validate our observation, the viral load in A549 cells were determined by infecting cells with GFP-tagged NDV by using flow cytometry, cells with negative control + NDV-GFP, m203 + NDV-GFP, and I203 + NDV-GFP samples. miR-203a-3p mimic treatment resulted in an increased NDV load, while the miR-203a-3p inhibitor (I203) led to a decrease in viral load (Fig. 5C-D). Similar results were observed in human peripheral blood mononuclear cells (hPBMCs) at 24 and 48-hours post-infection, further supporting the role of miR-203a-3p in enhancing the viral replication (Fig. 5E). We also investigated whether miR-203a-3p affects viral entry or restricts viral replication during the early stages of infection. We transfected A549 cells with either a negative control (NC) or miR-203a-3p mimic (m203). After 24 hours of transfection, the cells were infected with PR8, and samples were collected at 1, 3 and 12-hours post-infection. The viral load was quantified using qRT-PCR, which showed an increase in viral load at early time points post-infection. This indicates that miR-203a-3p does not interfere with viral entry or the early stages of infection (Fig. S5A). Altogether, these results suggest that overexpression of miR-203a-3p enhances viral replication by suppressing type I interferon genes, which are essential for antiviral defense. This finding provides valuable insight into the regulatory role of miR-203a-3p in host-pathogen interactions and highlights its potential as a therapeutic target for controlling viral infections.

**Figure 5:**
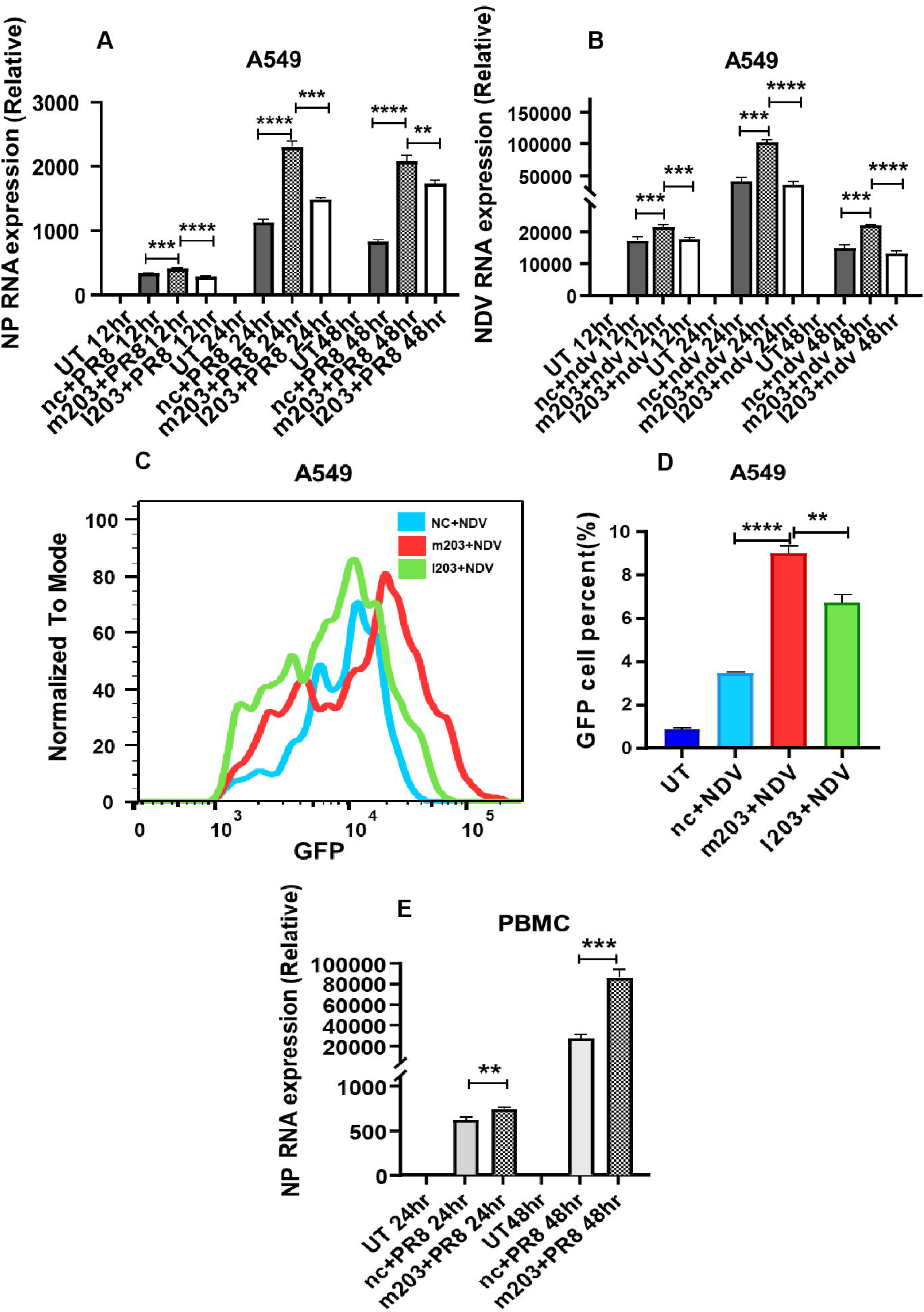
miR-203a-3p enhances RNA virus replication: (A-B) A549 cells were transfected with a miR-203a-3p mimic (m203a), a negative control (NC), or a miR-203a-3p inhibitor (I203) at a concentration of 50 nM. After 24 hours, cells were infected with Newcastle Disease Virus A/PR8/H1N1 (PR8) or (NDV) at a multiplicity of infection (MOI) of 1.5. Samples were collected at 12, 24, and 48 hours post-infection. (A) The relative expression of the NP gene was quantified by qRT-PCR. (B) The expression of the NDV L polymerase (L pol) gene was also measured at the same time points. (C-D) Fluorescence-activated cell sorting (FACS) analysis was performed to measure viral load. A549 cells were transfected with NC, m203a, or I203 at 50 nM. After 24 hours, cells were infected with NDV (MOI = 1.5) and incubated for an additional 24 hours. Cells were then trypsinized, washed with 1× PBS, fixed with 4% formaldehyde, and analyzed by flow cytometry for GFP. (E) A similar experiment was conducted using peripheral blood mononuclear cells (PBMCs) transfected with m203a or NC at 50 nM, followed by infection with A/PR8/H1N1. The relative expression of the NP gene was measured by qRT-PCR at 24 and 48 hours post-infection. Statistical significance is indicated as follows: * (p < 0.05), ** (p < 0.01), *** (p < 0.001), and****(p<0.0001).

## Discussion

The innate immune system relies on various sensors to detect viral components and initiate an immune response. These sensors include pattern recognition receptors (PRRs) such as Toll-like receptors (TLRs), retinoic acid-inducible gene I (RIG-I)-like receptors (RLRs), nucleotide-binding oligomerization domain (NOD)-like receptors (NLRs), and cytosolic DNA sensors(44, 45). Influenza virus triggers multiple signaling pathways inducing changes in immune and regulators of immune responses such as noncoding (small and long) RNAs. These alterations contribute to the development of antiviral immune response for viral clearance and development of immune homeostasis after viral clearance. Recent several report suggest that the pathogenic insult is also associated with cellular miRNA expression, and altered miRNA expression leads to the miRNA-mediated silencing of host and viral transcripts (46, 47). In this study, we found that the expression of miR-203a-3p was significantly elevated upon RNA virus infection, such as NDV and A/PR8/H1N1, as well as upon transfection with polyinosinic-polycytidylic acid (I:C)], a synthetic analog of viral double-stranded RNA that mimics viral infection.

The findings of this study highlight the crucial role of miR-203a-3p in regulation of type I interferon-mediated antiviral responses and promoting the replication of RNA viruses. By analyzing publicly available microarray datasets, we identified miR-203a-3p as a consistently upregulated microRNA during RNA virus infections. Bioinformatics analysis revealed that miR-203a-3p targets key components of the interferon signaling pathway, including members of the IFNA family, IFNAR1, JAK1, STAT1, and SOCS3. The same were also validated through luciferase reporter assays and site-directed mutagenesis in seed sequence of 3’UTR of the genes, demonstrating the specific binding of miR-203a-3p to the 3’ UTRs of these genes, which resulted in their downregulation.

The RNA immunoprecipitation (RIP) assay provided strong evidence for the direct interaction of miR-203a-3p with its target transcripts in association with Ago2, a vital component of the RNA-induced silencing complex (RISC). This interaction emphasizes the functional importance of miR-203a-3p in modulating the expression of genes involved in the type-I interferon pathway. Additionally, Western blot analysis supported these findings by showing decreased protein levels of JAK1 and IFNAR1 upon overexpression of miR-203a-3p, further confirming its regulatory effect. We explored the functional implications of miR-203a-3p in viral replication by assessing viral loads in cells via overexpression or inhibiting the endogenous miR-203a-3p. The results demonstrated that overexpression of miR-203a-3p enhances viral replication, as shown by increased levels of the A/PR8/H1N1 NP and NDV L- pol genes. Conversely, inhibiting miR-203a-3p led to reduced viral loads, underscoring its role as a positive regulator of viral replication. These observations were consistent across different cell lines and human primary cells, i.e., peripheral blood mononuclear cells (PBMCs), reinforcing the universality of this mechanism. Furthermore, the suppression of type I interferon signaling by miR-203a-3p was evidenced by decreased levels of IFN-β and IFN-α in cells transfected with miR-203a-3p mimics during viral infection. This suppression likely contributes to the observed increase in viral replication (Fig. 6), as type I interferons are essential for activating interferon-stimulated genes (ISGs) and mounting an effective antiviral response. Previously, it has been shown that miR-203a-3p is upregulated during influenza A virus (IAV) infection and suppresses H5N1 replication by targeting the down-regulator of transcription 1 (DR1). Another report showed that miR-203a-3p downregulates IFIT1/ISG56 expression(48, 49) The restoration of IFN-α and IFN-β expression following the inhibition of miR-203a-3p highlights its significance in antiviral immunity and immune homeostasis. Further research will deepen our understanding of miR-203a-3p and aid in developing innovative therapeutic strategies against RNA virus infections and complex autoimmune disease such as SLE.

**Figure 6:**
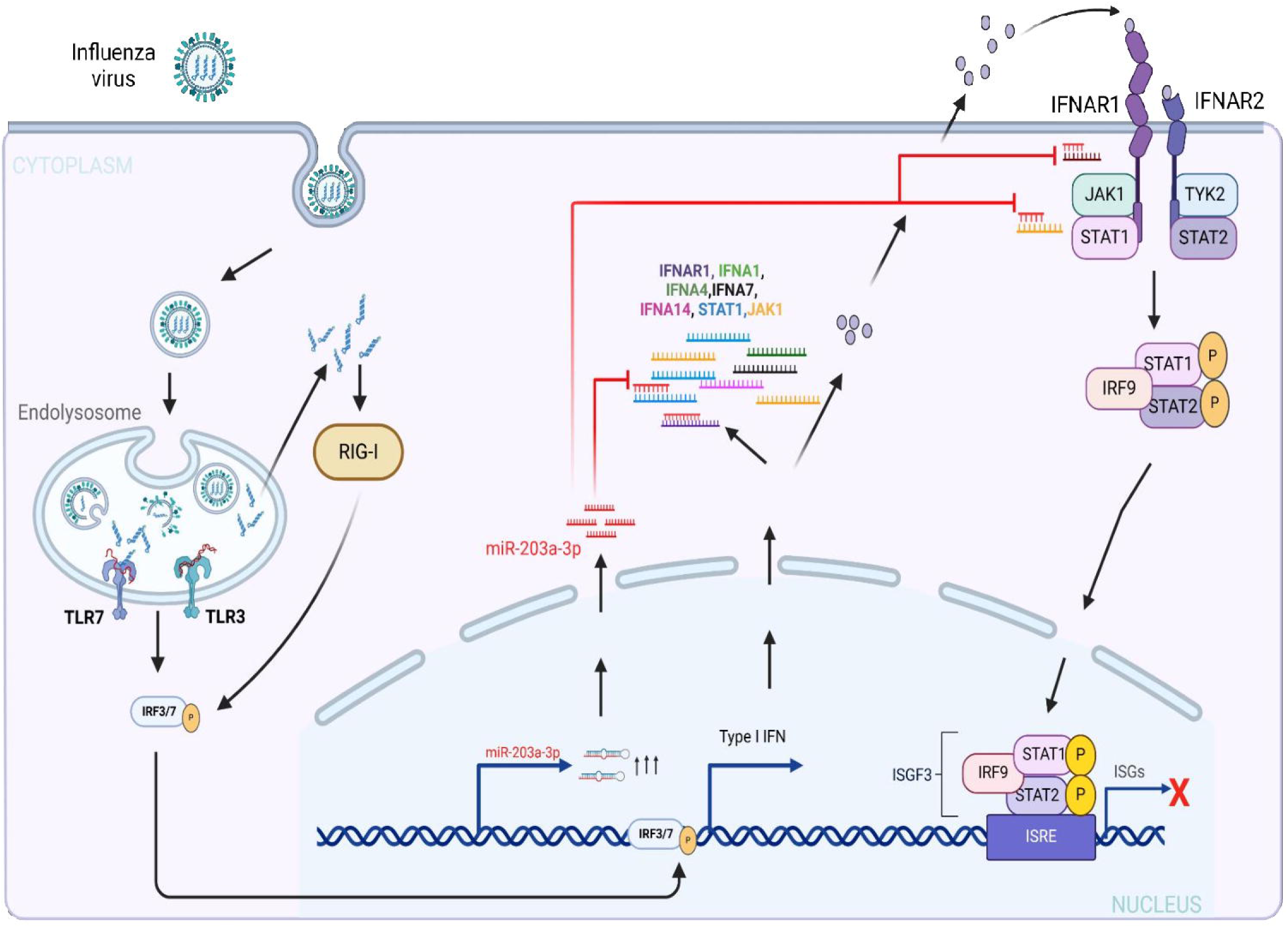
Proposed model of miR-203a-3p–mediated regulation of the Type I IFN response: The schematic illustrates that miR-203a-3p expression is induced following A/PR8/H1N1 and NDV infections. miR-203a-3p directly binds to the 3′ UTR of key molecules Type I IFN and type I IFN signaling pathway genes, leading to their downregulation of these genes. This suppression impairs the downstream antiviral activity of interferon-stimulated genes (ISGs), ultimately enhancing viral replication.

### Statistical analysis

All experiments were conducted with appropriate controls, including mock-transfected and untreated samples. Each experiment was independently replicated at least three times to ensure reproducibility. Statistical analyses were performed using GraphPad Prism software; the student’s unpaired two-tailed t-test was used for comparisons between two groups. Significance was determined at a p-value threshold of <0.05, ensuring the robustness and reliability of the findings.

## Acknowledgment

We sincerely thank R. Fouchier for providing the A/PR8/H1N1 reverse genetics system, P. Palese for the green fluorescent protein-expressing NDV (NDV-GFP), and IISER Bhopal for access to the Central Instrumentation Facility. We gratefully acknowledge the generous financial support from the DBT, New Delhi, and SERB, New Delhi. P.K. also thanks DBT for awarding the SRF.

## Author Contribution

P.K., A.K., Ak.K., and H.K. conceptualized the study. P.K. and A.K. performed experiments. A.K. analyzed data from the repository, and P.K. and A.K. did experiments on A/PR8/H1N11 and NDV in A549, 293T, HeLa, and PBMC cells. P.K., Ak.K., and H.K. prepared the manuscript; Kunal Arora helped with proofreading and editing, and H.K. supervised the entire project.

## Supplementary figures legends

**Figure S1: Pathway analysis of miR-203a-3p-targeted transcripts using Enrichr:** (A) BioPlanet pathway analysis reveals that miR-203a-3p targets key transcripts involved in xinterferon-alpha signaling, including the TRAF6-mediated IRF7 activation pathway. (B) This plot showing the gene pathways ranked based on their odds ratio and p-value (C) A549 cells were transfected with poly(I:C), and miR-203a-3p expression was assessed 24 hours post-transfection using qRT-PCR. Statistical significance is indicated as follows: * (p < 0.05), ** (p < 0.01), *** (p < 0.001), and **** (p < 0.0001).

**Figure S2: Pathway analysis of miR-378c and miR-199a-5p targeted transcripts using Enrichr:** Pathway analysis was performed using Enrichr to explore the targeted transcripts of miR-378c and miR-199a-5p. (A-B) The top 500 *miR-378c*–targeted transcripts were input into Enrichr, but no significant pathways related to the immune system were identified. (C-E) A similar analysis was conducted for *miR-199a-5p*, and no immune-related pathways were found in this case either.

**Figure S3: miR-203a-3p non-target controls:** (A) The IRF7 UTR does not contain any binding sites for miR-203a-3p; therefore, it was used as a non-target control in the luciferase assay. (B) IRF7 was also used in the RNA-induced silencing complex (RISC) assay, where it showed no significant interaction. Statistical significance is indicated as follows: * (p < 0.05), ** (p < 0.01), *** (p < 0.001), and **** (p < 0.0001).

**Figure S4: Viral load in interferon non-responsive cells:** Viral load was measured in interferon-deficient Vero E6 cells, which lack type I interferon receptors and do not respond to type I interferon signaling. Cells were infected with either A/PR8/H1N1 or NDV at a multiplicity of infection (MOI) of 1.5. Viral loads were quantified at 12 and 24 hours post-infection using qRT-PCR. (A) A/PR8/H1N1 viral load in Vero E6 cells was measured at 12 and 24 hours post-infection, while (B) NDV viral load was assessed at the same time points. Statistical significance is indicated as follows: * (p < 0.05), ** (p < 0.01), *** (p < 0.001), and **** (p < 0.0001).

**Figure S5: miR-203a-3p does not interfere with virus entry:** To determine whether miR- 203a-3p influences viral entry or restricts viral replication during the early stages of infection, A549 cells were transfected with either a negative control (NC) or miR-203a-3p mimic (m203). Twenty-four hours post-transfection, the cells were infected with A/PR8/H1N1 (PR8), and samples were collected at different time points (1, 3, and 12 hours). (A) PB1 expression levels were quantified using qRT-PCR. Statistical significance is indicated as follows: * (p < 0.05), ** (p < 0.01), *** (p < 0.001), and **** (p < 0.0001).

