## supplementary data for "Essential role of hsa-miR-203a-3p in type I Interferons immune homeostasis during Influenza and NDV infection"

**Fig. S1**

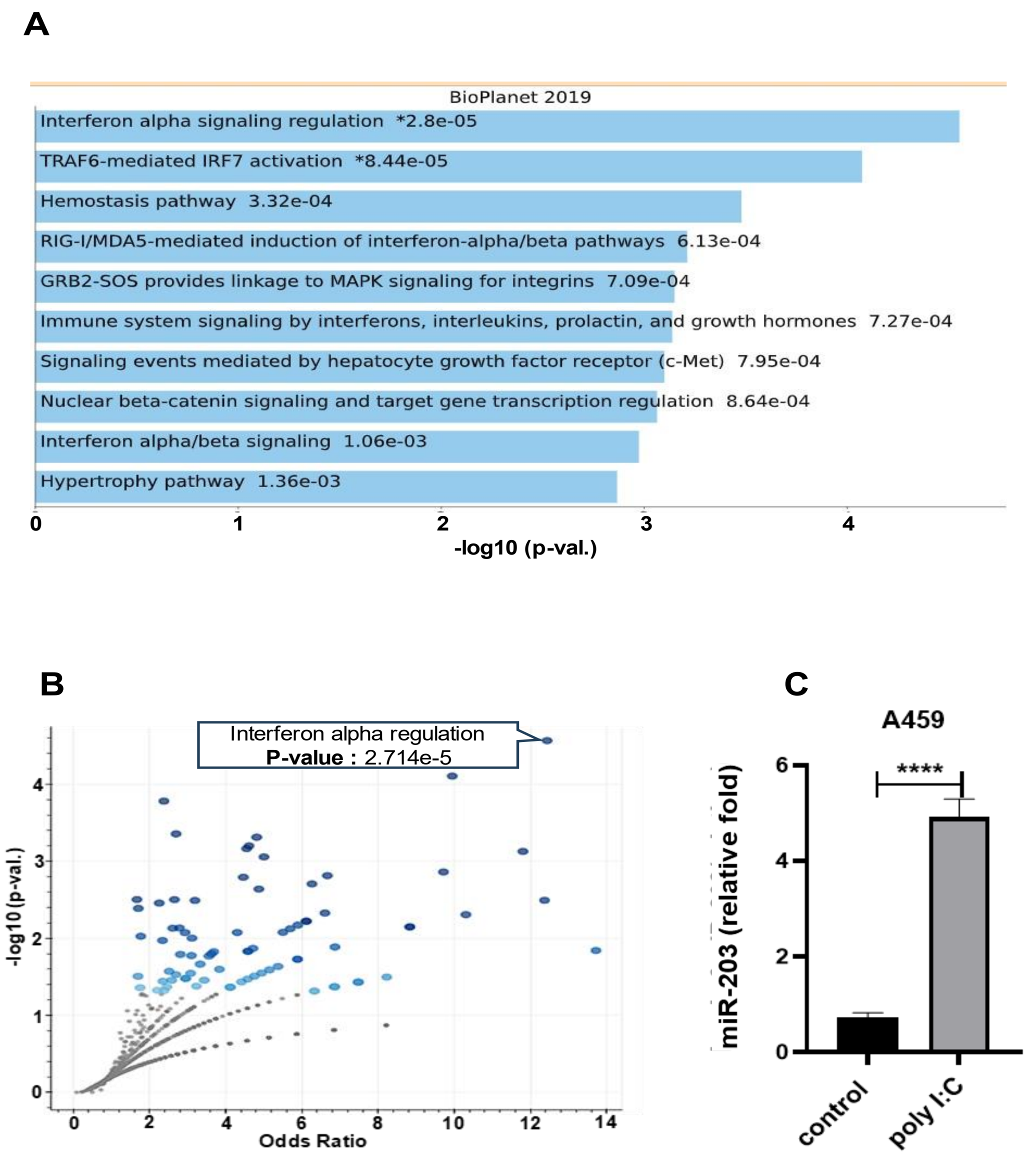


**Figure S1: Pathway analysis of miR-203a-3p targeted transcripts using Enrichr** (A) BioPlanet pathway analysis reveals that *miR-203a-3p* targets key transcripts involved in interferon-alpha signaling, including the TRAF6-mediated IRF7 activation pathway. (B) This plot showing the gene pathways ranked based on their odds ratio and p-value (C) A549 cells were transfected with poly(I:C), and *miR-203a-3p* expression was assessed 24 hours post-transfection using RT-PCR. Statistical significance is indicated as follows: * ($p < 0.05$), ** ($p < 0.01$), *** ($p < 0.001$), and **** ($p < 0.0001$).

Fig. S2

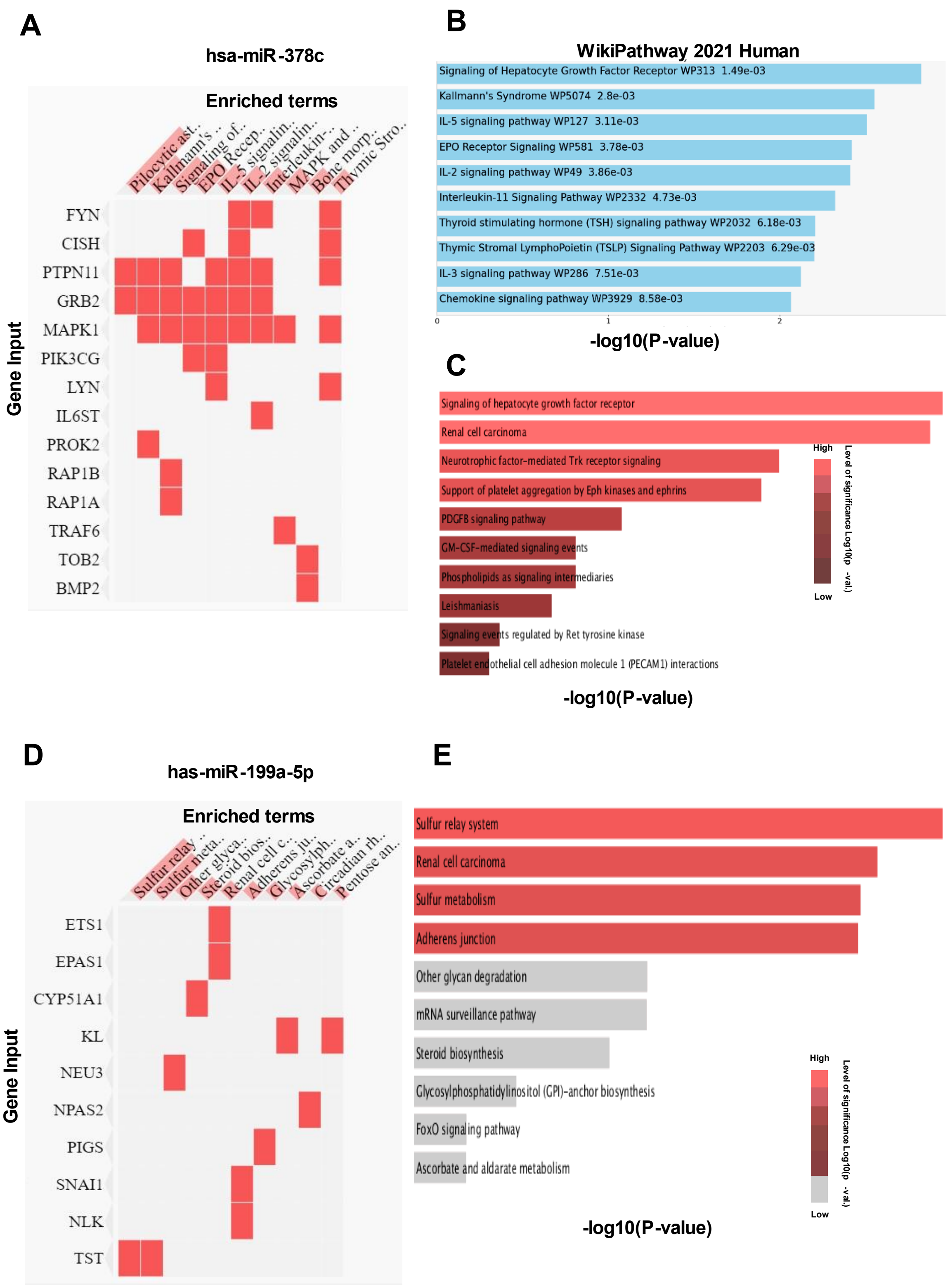
A
hsa-miR-378c
Enriched terms
Pilocytic ast..
Kallmann's ..
Signaling of..
EPO Recep..
IL-5 signalin..
IL-2 signalin..
Interleukin-..
MAPK and ..
Bone morp..
Thymic Stro..
Gene Input
FYN
CISH
PTPN11
GRB2
MAPK1
PIK3CG
LYN
IL6ST
PROK2
RAP1B
RAP1A
TRAF6
TOB2
BMP2
B
WikiPathway 2021 Human
Signaling of Hepatocyte Growth Factor Receptor WP313 1.49e-03
Kallmann's Syndrome WP5074 2.8e-03
IL-5 signaling pathway WP127 3.11e-03
EPO Receptor Signaling WP581 3.78e-03
IL-2 signaling pathway WP49 3.86e-03
Interleukin-11 Signaling Pathway WP2332 4.73e-03
Thyroid stimulating hormone (TSH) signaling pathway WP2032 6.18e-03
Thymic Stromal LymphoPoietin (TSLP) Signaling Pathway WP2203 6.29e-03
IL-3 signaling pathway WP286 7.51e-03
Chemokine signaling pathway WP3929 8.58e-03
0
1
2
-log10(P-value)
C
Signaling of hepatocyte growth factor receptor
Renal cell carcinoma
Neurotrophic factor-mediated Trk receptor signaling
Support of platelet aggregation by Eph kinases and ephrins
PDGFB signaling pathway
GM-CSF-mediated signaling events
Phospholipids as signaling intermediaries
Leishmaniasis
Signaling events regulated by Ret tyrosine kinase
Platelet endothelial cell adhesion molecule 1 (PECAM1) interactions
High
Low
Level of significance Log10(p -val.)
-log10(P-value)
D
has-miR-199a-5p
Enriched terms
Sulfur relay ..
Sulfur meta..
Other glyca..
Steroid bios..
Renal cell c..
Adherens ju..
Glycosylph..
Ascorbate a..
Circadian rh..
Pentose an..
Gene Input
ETS1
EPAS1
CYP51A1
KL
NEU3
NPAS2
PIGS
SNAI1
NLK
TST
E
Sulfur relay system
Renal cell carcinoma
Sulfur metabolism
Adherens junction
Other glycan degradation
mRNA surveillance pathway
Steroid biosynthesis
Glycosylphosphatidylinositol (GPI)-anchor biosynthesis
FoxO signaling pathway
Ascorbate and aldarate metabolism
High
Low
Level of significance Log10(p -val.)
-log10(P-value)

**Figure S2: Pathway analysis of miR-378c and miR-199a-5p targeted transcripts using Enrichr:** Pathway analysis was performed using Enrichr to explore the targeted transcripts of miR-378c and miR-199a-5p. (A-B) The top 500 *miR-378c*–targeted transcripts were input into Enrichr, but no significant pathways related to the immune system were identified. (C-E) A similar analysis was conducted for *miR-199a-5p*, and no immune-related pathways were found in this case either.

#### Fig. S3

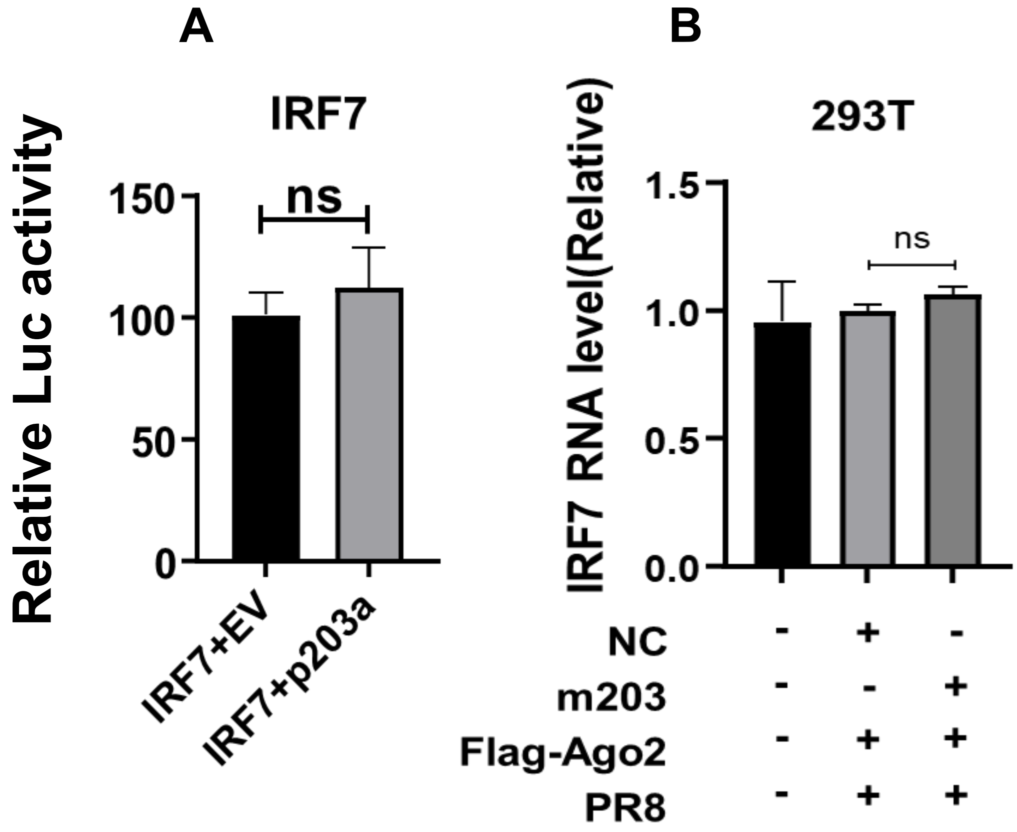


**Figure S3: miR-203a-3p non-target controls:** (A) The IRF7 UTR does not contain any binding sites for miR-203a-3p; therefore, it was used as a non-target control in the luciferase assay. (B) IRF7 was also used in the RNA-induced silencing complex (RISC) assay, where it showed no significant interaction. Statistical significance is indicated as follows: * ($p < 0.05$), ** ($p < 0.01$), *** ($p < 0.001$), and **** ($p < 0.0001$).

**Fig. S4**

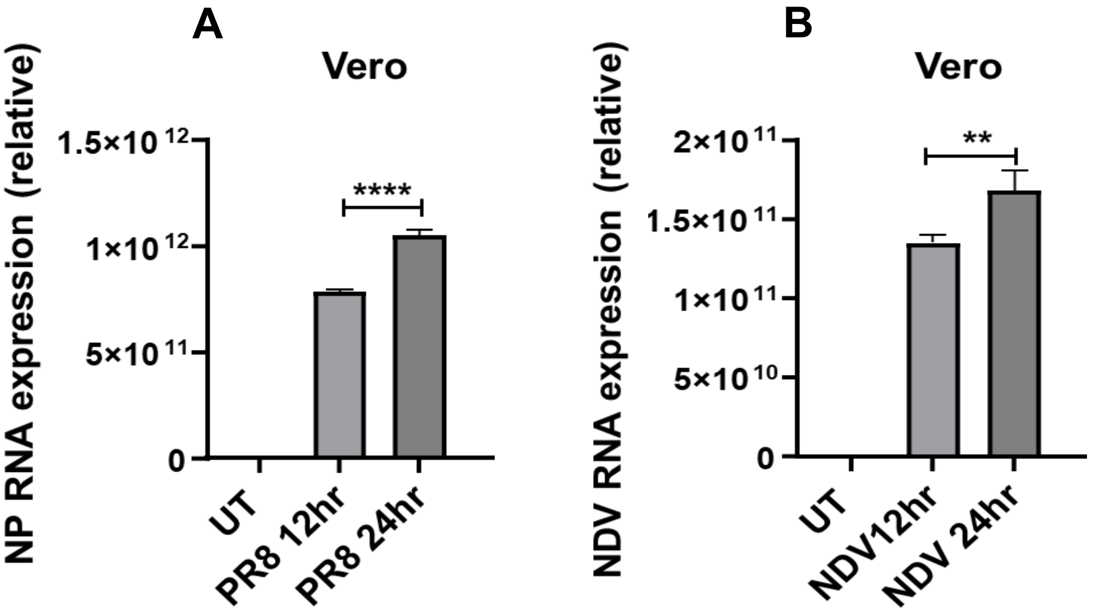


**Figure S4: Viral load in interferon non-responsive cells:** Viral load was measured in interferon-deficient Vero E6 cells, which lack type I interferon receptors and do not respond to type I interferon signaling. Cells were infected with either A/PR8/H1N1 or NDV at a multiplicity of infection (MOI) of 1.5. Viral loads were quantified at 12 and 24 hours post-infection using qRT-PCR. (A) A/PR8/H1N1 viral load in Vero E6 cells was measured at 12 and 24 hours post-infection, while (B) NDV viral load was assessed at the same time points. Statistical significance is indicated as follows: * ($p < 0.05$), ** ($p < 0.01$), *** ($p < 0.001$), and **** ($p < 0.0001$).

### Fig. S5

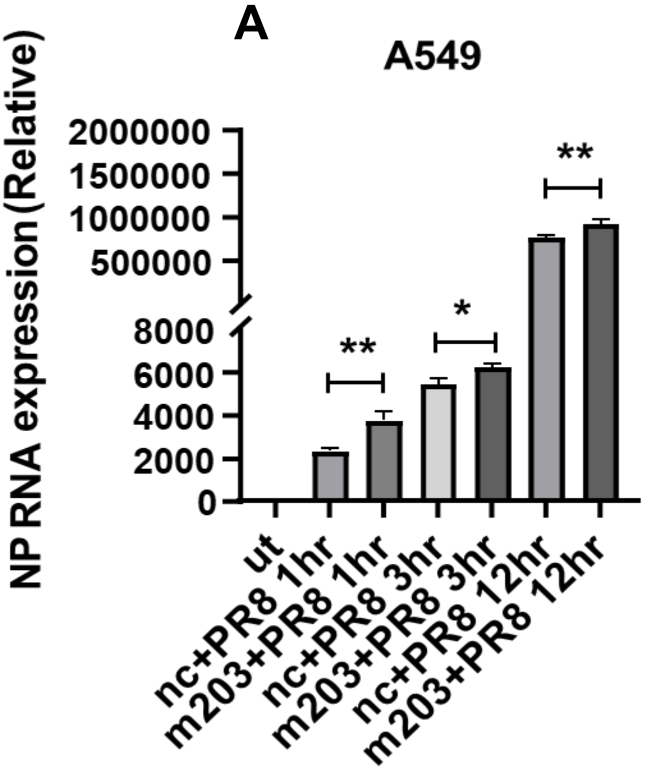


**Figure S5: miR-203a-3p does not interfere with virus entry:** To determine whether miR-203a-3p influences viral entry or restricts viral replication during the early stages of infection, A549 cells were transfected with either a negative control (NC) or miR-203a-3p mimic (m203). Twenty-four hours post-transfection, the cells were infected with A/PR8/H1N1 (PR8), and samples were collected at different time points (1, 3, and 12 hours). (A) PB1 expression levels were quantified using qRT-PCR. Statistical significance is indicated as follows: * ($p < 0.05$), ** ($p < 0.01$), *** ($p < 0.001$), and **** ($p < 0.0001$).
